# First-void urine and cervicovaginal brushes as sampling devices for vaginal microbiome studies: a proof-of-concept study

**DOI:** 10.1101/2025.03.14.643287

**Authors:** Maria Pinedo-Bardales, Anne Van Caesbroeck, Sarah Ahannach, Margo Bell, Severien Van Keer, Laura Téblick, Isabel Erreygers, Sam Bakelants, Sarah Van den Bosch, Sandra Condori-Catachura, Stijn Wittouck, Alex Vorsters, Sarah Lebeer

## Abstract

This proof-of-concept study compares first-void urine (FVU) and cervicovaginal brushes to vaginal swabs for microbiome profiling in ten women. FVU and brushes provided comparable results in most cases, though storage conditions influenced microbial composition, especially for FVU. Despite limitations in sample size and diversity in participants, findings support these methods’ potential for self-sampling in women’s health research, emphasizing the need for optimized techniques to improve vaginal microbiome profiling.

## Main section

The human vaginal microbiome is a cornerstone of female reproductive health, influencing susceptibility to vaginal conditions, pregnancy complications, and sexually transmitted and urinary infections^1^. Understanding this microbiome is critical for advancing preventive, diagnostic and therapeutic interventions, yet its intimate and dynamic nature poses challenges for accurate, representative and more inclusive sampling methods.

Current approaches for mapping and assessing the vaginal microbiome predominantly rely on direct sampling through vaginal swabs^2,3^ or more invasive techniques such as lavages^4,5^. The large-scale citizen-science project Isala^2^ (https://isala.be/en/) and its international Isala sisterhood^6^ leverage self-collected vaginal swabs to study the vaginal microbiome. Initially launched in Flanders (Belgium), Isala analyzed samples from more than 3,345 women and has since expanded globally, involving thousands of women from countries in America, Africa and Europe^6^. While vaginal swabs have proven effective for microbiome profiling^7^, their use can be perceived as uncomfortable by some women. Cultural norms, age, stigmas, and privacy concerns in certain communities can also limit their acceptability and accessibility^8^.

To address these issues, first-void urine (FVU) samples offer a non-invasive, user-friendly alternative for collecting intimate samples. FVU refers to the initial stream of urine that flushes away cervicovaginal secretions that accumulated between the small labia and the urethral opening^9^. This sample may be collected at any time of the day^10^, and allows for the detection of bacteria, viral particles, DNA, methylation markers, proteins, metabolites, and immunological biomarkers including cytokines, antibodies and immune cells^11–16^. In addition, while once considered sterile in healthy individuals, the urinary tract is now recognized as a diverse microbial environment, even in the absence of infection, with *Lactobacillus, Prevotella* and *Gardnerella* among the most frequently reported genera in non-FVU urine samples from healthy women^17,18^.

Another promising approach for intimate microbiome profiling is the use of cervicovaginal brushes. These devices allow to obtain more cells from the vaginal wall, offering valuable insights into vaginal microbial communities and their metatranscriptomic activity, as well as host-immune interactions^19–22^. Designed for self-collection, they also enhance participant comfort and compliance, making them particularly useful for large-scale studies^23–25^.

This proof-of-concept study aimed to evaluate the effectiveness of FVU samples and cervicovaginal brushes as self-sampling devices for vaginal microbiome profiling relative to vaginal swabs. Ten healthy reproductive-age women were enrolled in Antwerp (Belgium) after ethical approval (B300201734129). Participants were equally divided into two groups and self-collected four samples each (Figure 1a). FVU samples were collected using a 20 mL Colli-Pee® device (DNA Genotek Inc, Ottawa, Canada), prefilled with 1:3 urine conservation medium (UCM) (DNA Genotek Inc, Ottawa Canada). For collection of vaginal swabs, eNAT® swabs (Copan, Brescia, Italy) were used. These devices contain 1 mL of a lysis buffer designed for prolonged stabilization and preservation of microbial DNA. Cervicovaginal samples were self-collected using Evalyn® cervicovaginal brushes (Rovers Medical Devices, Oss, The Netherlands), which were resuspended in 20 mL ThinPrep PreservCyt solution (Hologic, Massachusetts, USA). For both FVU samples and vaginal swabs, two aliquots were obtained to evaluate two different storage conditions per device (Figure 1b) for their impact on sample quality, as determined by bacterial and human DNA content. A 1 mL aliquot of each diluted sample was used for DNA extraction.

**Figure 1:**
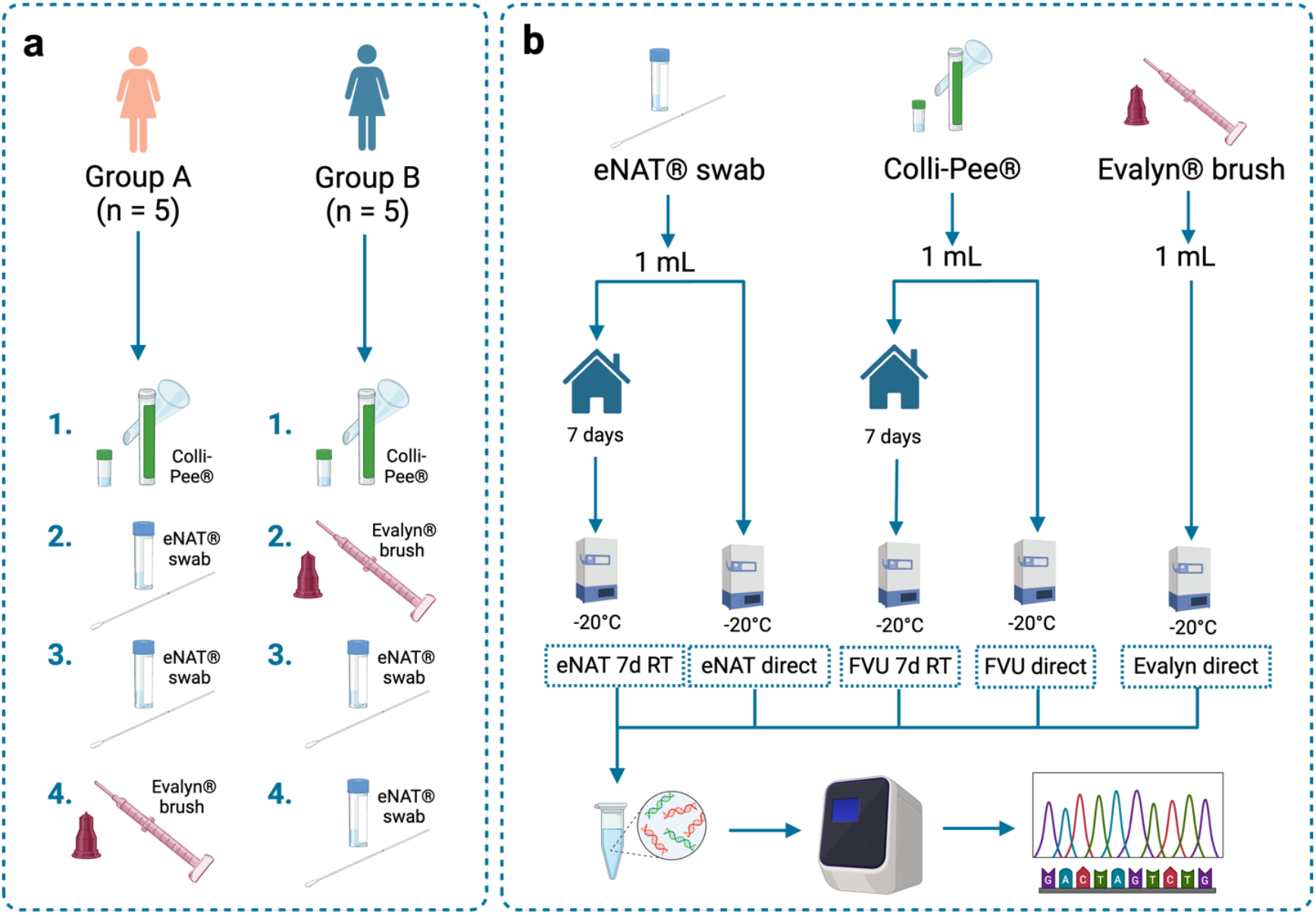
Overview of the study workflow. **a)** Sampling sequence for each group, outlining the order in which participants collected first-void urine samples (FVU; cfr. Colli-Pee®), vaginal swabs (cfr. eNAT® swab), and cervicovaginal brushes (cfr. Evalyn® brush). **b)** Comparison of two storage conditions tested for FVU samples and vaginal swabs (seven days at RT versus directly frozen), followed by downstream analytical processes. RT = room temperature.

After DNA extraction, samples showed human DNA concentrations ranging from 251 ng/μL to 3.16 x 10^6^ ng/μL (Supplementary Fig. 1). When assessing bacterial DNA (Figure 2a), eNAT® swabs demonstrated a significantly higher bacterial DNA concentration (averaging 2.20 x 10^9^ ng/μL) compared to samples collected using the Colli-Pee® device (p=0.0021) and the Evalyn® brush (p=0.00032). This difference may be due to variations in dilution across the buffers of each sampling device. For instance, eNAT® swabs contained 1 mL of lysis buffer, Colli-Pee® devices were pre-filled with 6.66 mL of UCM, and Evalyn® brushes were resuspended in 20 mL of ThinPrep PreservCyt solution. We also assessed the impact of storing of the samples and found that storage for seven days at room temperature (RT) did not impact bacterial or human DNA concentrations for either the eNAT® swabs or urine samples (Supplementary Fig. 1 and Fig. 2a). Regarding the bacterial-to-human DNA ratio, which is a key metric for 16S rRNA amplicon sequencing, the eNAT® swab showed a significantly higher value, particularly when compared to Colli-Pee® samples, including aliquots stored at RT (Figure 2b).

**Figure 2:**
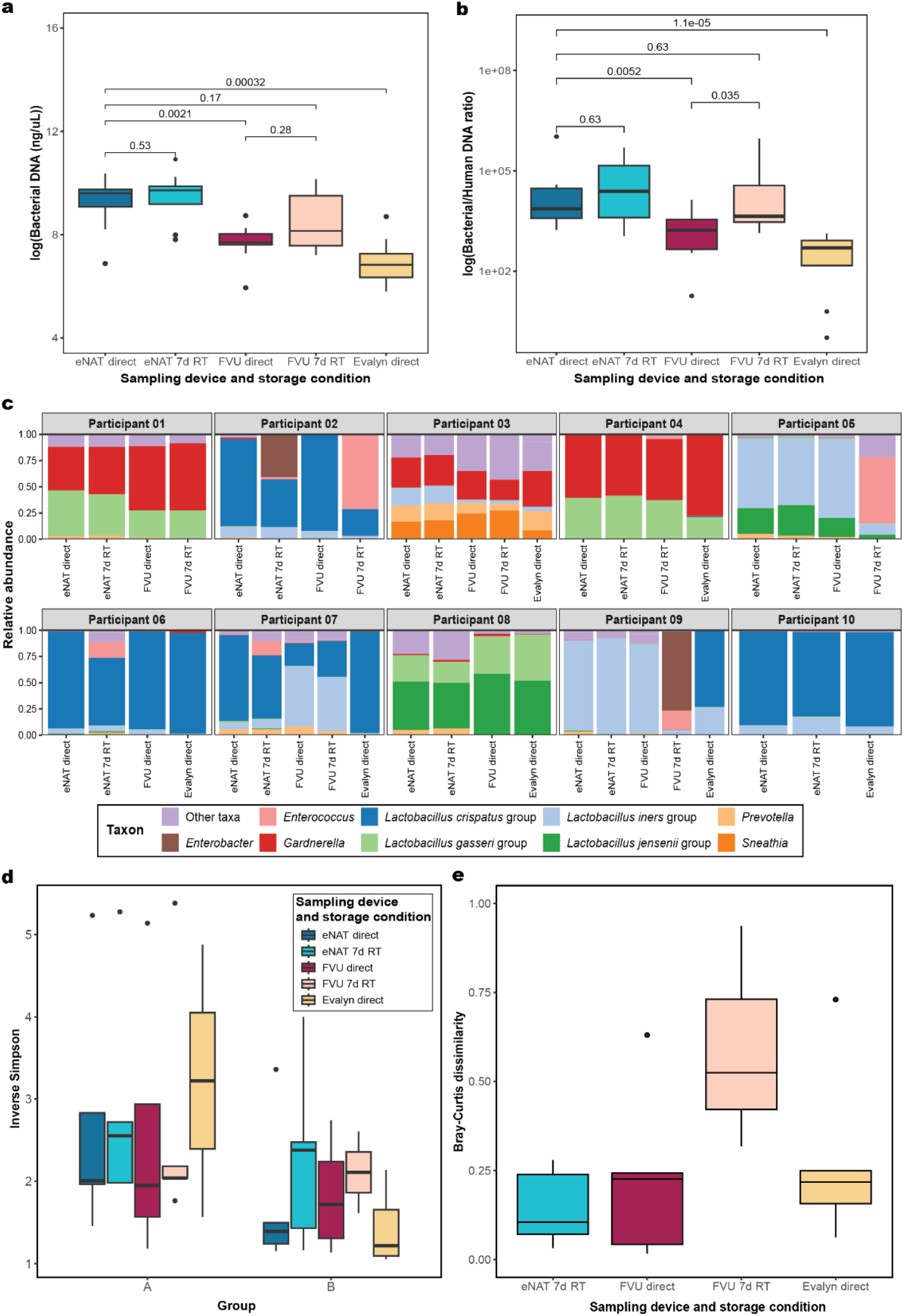
Overview of the comparison of the different sampling devices and storage conditions in terms of bacterial DNA concentration (a), bacterial-to-human DNA ratio (b), taxonomic composition (c), alpha diversity (d) and beta diversity (e).

We then used dual-index pair-end sequencing for amplification of the V4 region of the 16S rRNA gene and first, analyzed the sequencing reads and taxonomic profiles of the negative controls (Supplementary Fig. 2a and Fig. 2b). On average, 228 reads were obtained from the controls, with a maximum DNA concentration (5.49 ng/μL) in the control of the ThinPrep solution, though no bacterial reads were detected in this sample. The highest read concentration was in the second DNA extraction control, with *Gardnerella* and *Lactobacillus gasseri* being the taxa with the highest relative abundance on this sample.

After quality control of sequencing reads based on the controls, FVU samples stored for seven days at RT yielded the highest mean total read count (29,007 reads), followed by FVU directly stored (20,602 reads) and eNAT® directly stored (20,235 reads) (Supplementary Fig. 2a). We then examined the distribution of high-quality reads per participant through microbiome profiling. Taxonomic profiles were successfully obtained for all samples of three participants (participant 03, 07 and 09; Figure 2c). In addition to eNAT® swabs, FVU samples directly stored performed well, with eight out of ten participants having high-quality reads, confirming their reliability under optimal storage conditions. In contrast, high-quality reads were undetectable in seven out of ten samples collected with the Evalyn® brush or in FVU samples stored for seven days at RT. This may be attributed to factors such as suboptimal DNA preservation under certain storage conditions, or the inherently low microbial biomass in some sampling devices^26,27^.

We then also analyzed the taxonomic composition of the samples. Based on the eNAT® swabs, the vaginal microbiome of nine out of ten participants were dominated (relative abundance of 30% or higher) by lactobacilli, primarily from the *Lactobacillus crispatus* group (Figure 2c). One participant exhibited a more diverse profile, including species belonging to *Sneathia, Prevotella, Lactobacillus iners* group and *Gardnerella* genera. In seven out of ten participants, the taxonomic composition remained stable even after storing the swabs or the FVU samples for seven days at RT. However, storage led to an increased presence of usually less abundant vaginal taxa, such as *Enterococcus* and *Enterobacter*, possibly due to microbial shifts under suboptimal storage conditions or overgrowth of facultative anaerobes at RT. Furthermore, we cannot exclude the possibility of contamination with bacteria originating from another niche such as with fecal or skin bacteria. In other studies, microbial biomass in FVU also closely resembles vaginal microbiota in optimal conditions^28–30^.

To evaluate a possible taxonomic bias influenced by the sampling device, we analyzed the alpha diversity measured using the inverse Simpson per different group (Figure 2c). Higher dissimilarities were observed in Group A, where the sampling order was as follows: first the FVU sample, followed by two vaginal eNAT® swabs, and finally the Evalyn® brush; compared to Group B where the order was FVU, followed by the Evalyn® brush, and finally the two vaginal eNAT® swabs. When assessing the impact of storage at RT on microbiome composition, the Bray-Curtis dissimilarity index (Figure 2d) indicated lower dissimilarity between taxonomic profiles obtained from FVU samples and cervicovaginal brushes that were directly frozen after collection compared to those obtained from the eNAT® swabs for each participant. In contrast, higher dissimilarity was observed for profiles from FVU samples stored for seven days at RT.

Despite our findings, some limitations are inherent to this proof-of-concept study. The small sample size limits robust comparisons, and variations in sample collection, storage and handling may have introduced biases. High-quality reads were lacking in some samples, particularly from cervicovaginal brushes, affecting data comparability. However, the consistent taxonomic similarity between the profiles from vaginal swabs and FVU samples directly frozen, along with certain vaginal brushes, highlights their potential for vaginal microbiome research. From a citizen-science perspective, ease of sample collection is key^31^. FVU samples, being non-invasive and simpler to collect compared to vaginal swabs or brushes, represent a practical potential proxy for assessing vaginal microbiome in large-scale or remote studies, or in projects including hard-to-reach populations such as girls under 18 years old or other women uncomfortable with self-sampling.

Overall, this proof-of-concept study provides valuable insights into vaginal microbiome profiling using FVU samples and cervicovaginal brushes compared to vaginal swabs. Notably, eNAT® swabs yielded higher bacterial content, likely explained by the differences in the dilution buffers of each sampling device. Storage conditions also influenced microbial composition, with FVU samples stored for seven days at RT showing greater dissimilarity compared to those directly frozen after collection. Moreover, this proof-of-concept study highlights how other self-sampling devices can be used in vaginal microbiome research^31^, enabling more accessible and scalable methods for microbiome composition analysis. Such improvements could strengthen our understanding of the vaginal microbiota and their role in women’s health, paving the way for large-scale applications in future research projects.

## Methods

### Ethics approval

This study was approved by the Institutional Review Board (or Ethics Committee) of UZA/University of Antwerp (B300201734129) and conducted according to the guidelines of the Declaration of Helsinki. Written informed consent was appropriately obtained from all participants.

### Participants recruitment and sample collection

Ten self-reported healthy, reproductive-age women were recruited at the Centre for Evaluation of Vaccination of the University of Antwerp (Antwerp, Belgium) in October 2023. Sample collection was conducted in two groups of five participants, each following a different sampling order (Fig. 1a). All samples were self-collected.

Group A participants collected a FVU sample using the 20 mL Colli-Pee® device (DNA Genotek Inc, Ottawa, Canada), prefilled with 1:3 urine conservation medium (UCM) (DNA Genotek Inc, Ottawa, Canada). Participants were instructed not to wash their genitals thoroughly and to avoid urinating for at least 1 hour before collection. This was followed by the collection of two vaginal samples using eNAT® swabs (Copan, Brescia, Italy), containing 1 mL of a lysis buffer, and a cervicovaginal sample using the Evalyn® brush (Rovers Medical Services B.V., Oss, The Netherlands). In Group B, participants collected samples using the same devices, but in a different order: first the FVU sample, followed by the cervicovaginal brush and then two vaginal swabs. The Evalyn® brushes were resuspended in 20 mL of ThinPrep PreservCyt solution (Hologic, Massachusetts, USA). All samples were aliquoted in 1 mL Eppendorf tubes after thorough vortexing for 15-30 seconds. One of each participant’s vaginal swabs, and one 1 mL aliquot of FVU were kept at RT for up to seven days and subsequently frozen at -20°C. All other aliquots were immediately frozen at -20°C.

### DNA extraction and 16S rDNA amplicon sequencing

DNA was extracted from all 1 mL aliquots using the DNeasy PowerSoil Pro Kit (Qiagen, Germany) following the procedure described by Ahannach *et al*.^32^. The V4 region of the 16S rRNA gene of each bacterial DNA was amplified using standard primers altered for dual-index pair-end sequencing^33^. Library preparation was performed using standard in-house conditions^2^. Dual-index paired-end sequencing was performed on a MiSeq Desktop sequencer (Illumina). The sequencing run encompassed all DNA samples alongside controls for the UCM of the Colli-Pee device (DNA Genotek Inc, Ottawa Canada), ThinPrep PreservCyt solution, the DNA extraction processes and polymerase chain reaction (PCR) using PCR-grade water. For the latter two type of controls mentioned, two controls from each were considered.

### Quantitative PCR (qPCR)

qPCR was used to quantify both absolute bacterial and human DNA concentrations in samples after DNA extraction. The analysis was conducted in duplicates using the StepOnePlus Real-Time PCR System (v.2.0; Applied Biosystems, Foster City, California, USA), SYBR Green chemistry (PowerUp SYBR Green Master Mix, Applied Biosystems, Foster City, California, USA). The human housekeeping gene peptidylprolyl isomerase (PPIA)^34^ and universal bacterial 16S rRNA primers (338F and 518R)^33^ were used to estimate the human and bacterial DNA concentrations, respectively, using the standard curves previously described by Ahannach *et al*.^32^

### Bioinformatic analyses

Quality control and processing of amplicon reads were performed with the R package DADA2 version 1.22.0^35^ following previously established pipelines detailed by Lebeer *et al*.^2^. The merged and denoised reads (ASVs) underwent taxonomic annotation from the phylum to the genus level using the assignTaxonomy function and the Genome Taxonomy Database (GTDB)^36^ ssu archive (version 207). To enhance taxonomic resolution specifically for the genus *Lactobacillus*, ASVs from this genus were reclassified into nine different subgenera validated previously in vaginal samples from the Isala cohort^2^.

Taxon and sample quality control measures were applied as follows. ASVs identified as non-bacterial, including mitochondria, and those exceeding 260 bases in length were excluded. Samples were filtered based on two criteria: first, a DNA concentration above the maximum value of a negative control determined by manual inspection (5.49 ng/μL, Supplementary fig. 2a); and secondly, a read concentration (total read count divided by volume) above the maximum value of a negative control (113.2 reads/μL), determined by manual inspection (Supplementary Figure 2a). Taxonomic profiles of high-quality samples and diversity metrics were visualized using the in-house developed R package tidytacos (https://github.com/LebeerLab/tidytacos)^37^. The Wilcoxon test was used to compare the concentrations of the sampling device and protocol tested.

## Acknowledgements

We thank all the volunteers who participated in this study and Nele Van de Vliet for her help in processing the samples. The authors acknowledge the European Research Council (starting grant Lacto-Be 852600 of S.L and starting grant URISAMP 101040588 of A.V.), the Special Research Fund of the University of Antwerp (TT(ZAP)BOF 48897 of A.V.C.), the Research Foundation—Flanders (FWO; FWO Research project G049022N of S.L., FWO SBO Devenir project S006426, doctoral grant 11PJK4N of M.B., postdoctoral grant 12AZ624N of S.W.), and VLIR-UOS (project PE2022TEA519A102, for S.C.C and S.L.).

## Author contributions

A.V.C., S.A., M.B, S.V.K., L.T., A.V. and S.L. designed the study and worked on the conceptualization of the research project. A.V.C., M.P.B., S.A., M.B., I.E. and S.V.B. carried out the experimental and logistical work. A.V.C., I.E. and S.B. were responsible for the biobanking of all collected samples. M.P.B. and S.W. processed the sequencing data and performed the biostatistical analyses. M.P.B., S.A. and S.V.B. worked on the visualizations. M.P.B., A.V.C., S.A., M.B., I.E., S.V.B., S.C.C., S.W., A.V. and S.L. contributed to the interpretation of the results. M.P.B., S.A., A.V.C., M.B., I.E., A.V. and S.L wrote the original manuscript. All authors contributed to reviewing and editing of the final manuscript.

