## Supplementary Figures for "First-void urine and cervicovaginal brushes as sampling devices for vaginal microbiome studies: a proof-of-concept study"

* shared first author

$ shared senior author

° corresponding author

### Supplementary Figures

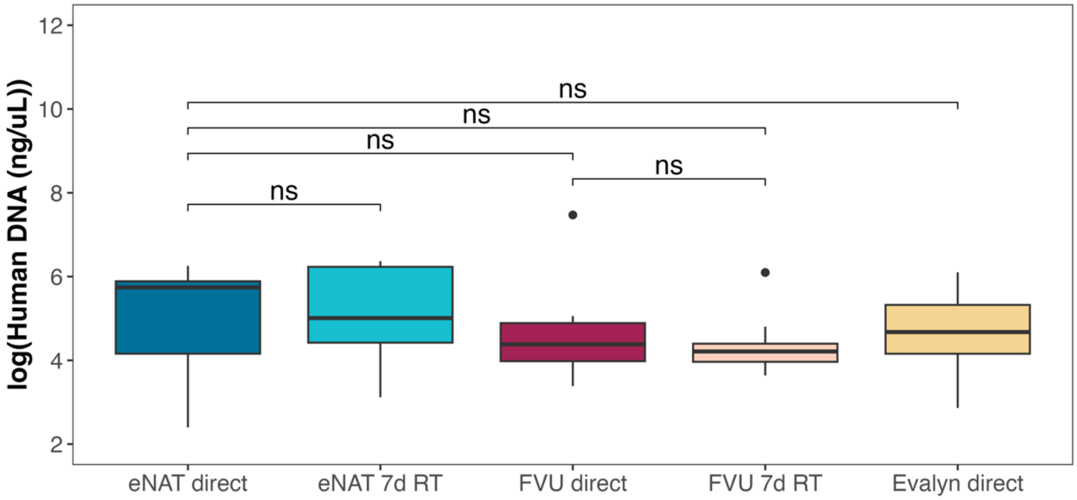


**Supplementary Figure 1: Human DNA concentration in the samples after DNA extraction.** Human DNA concentration was measured using qPCR including biological replicates. ns = No significance; eNAT direct = eNAT swab frozen immediately; eNAT 7d RT = eNAT swabs stored for seven days at room temperature (RT); FVU = First-void urine samples frozen immediately; FVU 7d RT = First-void urine samples stored for seven days at room temperature

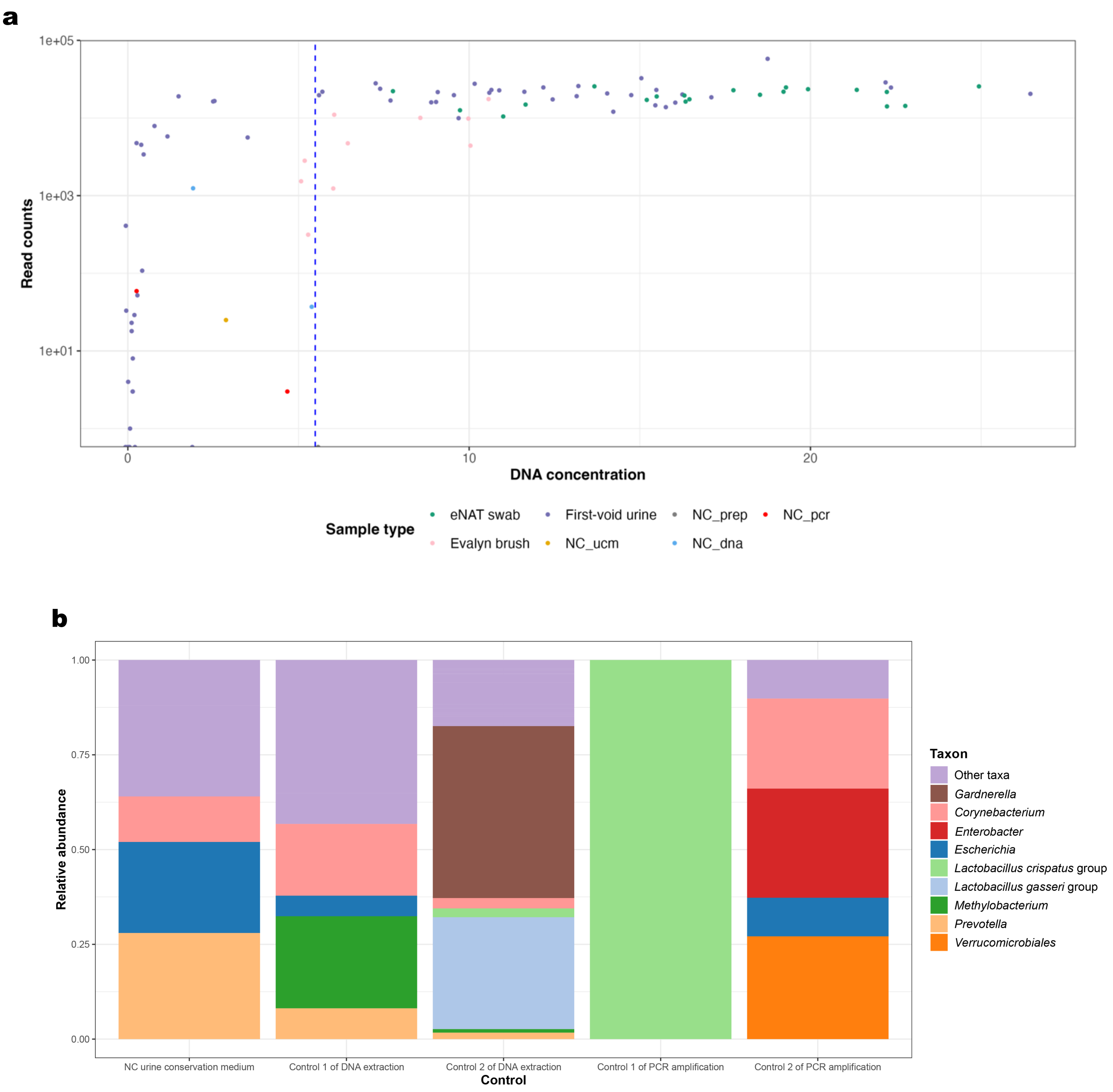


**Supplementary Figure 2: Quality control of the samples based on the negative controls.** The sequencing run encompassed all DNA samples alongside controls. Read concentration and DNA concentration for both the samples and controls are presented in Figure 2a, while the taxonomic composition of the controls is shown in Figure 2b. NC_ucm = Negative control (NC) of the urine conservation medium. NC_prep = NC of the ThinPrep PreservCyt solution. NC_dna = DNA extraction control. NC_pcr = PCR cleanup controls.
